# Liana cutting accelerates tropical forest recovery at a fraction of the cost of tree planting

**DOI:** 10.64898/2026.03.02.709120

**Authors:** Toby D. Jackson, Lucy V. J Beese, Andy Hector, Eleanor E. Jackson, Michael O’Brien, Gianluca Cerullo, David A. Coomes, David F. R. P. Burslem, Fabian J. Fischer, Christopher D. Philipson, Elia Godoong, Charissa Wong, Martin Svátek, Michele Dalponte, Wan Shafrina Wan Mohd Jaafar, Tommaso Jucker

**Affiliations:** School of Biological Sciences, University of Bristol, Bristol, UK; Department of Biology & Leverhulme Centre for Nature Recovery, University of Oxford, UK; Estación Experimental de Zonas Áridas, Consejo Superior de Investigaciones Científicas, Almería, Spain; Southeast Asia Rainforest Research Partnership (SEARRP), Kota Kinabalu, Sabah, Malaysia; Department of Forest Ecosystems and Society, Oregon State University, Corvallis, OR, USA; Department of Zoology, University of Cambridge, Cambridge, UK; Conservation Research Institute, University of Cambridge, Cambridge, UK; Department of Plant Sciences, University of Cambridge, UK; School of Biological Sciences, University of Aberdeen, Cruickshank Building, Aberdeen, AB24 3UU, UK; Ecosystem Dynamics and Forest Management Group, School of Life Sciences, Technische Universität München, Freising, Germany; belian.earth, York Eco Business Centre (Office 12), Amy Johnson Way, Clifton Moor, York, England, YO30 4AG; Faculty of Tropical Forestry, Universiti Malaysia Sabah; Forest Research Centre, Sabah Forestry Department, Sandakan, Sabah, Malaysia; Department of Forest Botany, Dendrology and Geobiocoenology, Faculty of Forestry and Wood Technology, Mendel University in Brno, Brno, Czech Republic; Research and Innovation Centre, Edmund Mach Foundation, San Michele all’Adige (TN), Italy; Earth Observation Centre, Institute of Climate Change, National University of Malaysia (UKM), 43600 Bangi, Selangor, Malaysia; Department of Geographical Sciences, University of Maryland, 2181 Samuel J. LeFrak Hall, 7251 Preinkert Drive, College Park, MD, USA

## Abstract

We urgently need to restore degraded tropical forests to mitigate the climate and biodiversity crises, but how to do so rapidly and cost-effectively remains an open question. Here we provide a long-term, landscape-scale assessment of the effectiveness of enrichment tree planting and liana cutting, the two most common restoration interventions used across many tropical regions. Leveraging one of the world’s largest and longest running forest restoration experiments, we used repeat airborne laser scanning to track the 3D structural recovery of 500 ha of selectively logged rainforest in Borneo. Over an 18-year period, enrichment planting increased mean canopy height by 1.6 m relative to unplanted controls. Remarkably, liana cutting increased canopy height more than four times faster (3.7 m over just 9 years). This recovery was jointly driven by accelerated canopy gap closure, enhanced tree growth, and a 50% reduction in tree mortality. Given that liana cutting is around 10 times cheaper to implement than enrichment planting, our results suggest it provides a cost-effective, scalable solution to accelerate the structural recovery of logged tropical forests.

## Introduction

Over the past century, vast portions of the world’s tropical forests have been fragmented and selectively logged, leaving more than half of all remaining tropical forests in a degraded state^1^. Restoring these degraded forests is essential to meeting international climate goals, such as limiting global warming to 2°C, and preventing further biodiversity loss^2,3^. Indeed, tropical forest restoration is widely recognized as one of the most impactful actions we can take to slow climate change^4,5^, and is central to global initiatives like the UN Decade on Ecosystem Restoration. A major threat to the recovery of these degraded forests are lianas (woody vines), which often proliferate after disturbance and can outcompete trees for light, water and nutrients^6^. Cutting lianas is therefore increasingly seen as a promising way to restore degraded tropical forests, especially as it is relatively cheap to implement^7^. However, while its effectiveness has been demonstrated in Central and South America^8–10^, it is unclear whether liana cutting will be as effective in other tropical regions such as Southeast Asia, where other restoration interventions such as enrichment tree planting are widely used.

The tropical forests of Southeast Asia are unique in that a significant proportion of their largest trees belong to a single family, the Dipterocarpaceae. These dipterocarp trees, which can reach heights of up to 100 meters in Borneo^11^, contribute substantially to the high carbon density of these forests^12^. Selective logging in Borneo generally targets large dipterocarps for their timber, thus altering forest structure and composition. However, removing these mature trees dramatically reduces seed production, which puts these forests at risk of regeneration failure^13^. This risk is particularly acute because dipterocarp seeds do not survive in the soil, meaning that there is no seed bank once mature trees have been removed. For this reason, restoration of selectively logged forests in Borneo often focuses on enrichment planting with dipterocarp seedlings^14,15^. However, enrichment planting is expensive ($1500-2500 ha^-1^), and many seedlings die in the first few years after planting^15^. Furthermore, enrichment planting and liana cutting are likely to have distinct impacts on forest structural recovery, with implications not just for carbon storage but also habitat complexity and biodiversity^16^. Unfortunately, we currently lack any quantitative comparison of their effects. Given that financial resources for restoration are limited, we urgently need to understand which approaches work best for Borneo’s rainforests.

To address this major knowledge gap, we acquired repeat airborne laser scanning (ALS) data over one of the world’s largest and longest-running tropical forest restoration experiments: the Sabah Biodiversity Experiment (SBE). SBE was established in 2002 and covers 500 hectares of selectively logged dipterocarp forest in Malaysian Borneo^17^. The experimental design of SBE allowed us to compare the impacts of enrichment planting and liana cutting against control plots with no intervention after selective logging, as well as benchmarking these against a nearby primary forest at Danum Valley. Using the repeat ALS data, we tracked how key attributes of canopy 3D structure recovered over time across each of these restoration treatments. This helped us disentangle different components of forest recovery following restoration, distinguishing between understory growth, canopy gap recovery, canopy growth and disturbances due to tree mortality and branch falls.

## Results

### Liana cutting is cost-effective in Borneo

In 2020, 18 years after the enrichment planting started, planted plots were 1.6 m ±0.7 m taller (mean ±standard error) than the control plots (p = 0.02), accounting for the effects of elevation (S1). At the same point, 9 years after treatment started, the liana cutting plots were 3.7 ±0.9 m taller than the control plots (p < 0.01), and 2.1 ±0.6 m taller than the enrichment planting plots (p < 0.01). Previous work in Sabah^12^ indicates that an increase in mean canopy height of 1 m represents approximately 5.2 Mg of additional aboveground carbon storage per hectare (S2). Liana cutting therefore led to an additional gain of +1.31 ±0.13 Mg C ha^-1^ yr^-1^, three times greater than that of enrichment planting (+0.38 ±0.03 Mg C ha^-1^ yr^-1^).

Liana cutting is also substantially cheaper to implement than enrichment planting. In Malaysia, liana cutting costs approximately USD$140-330 per hectare, while enrichment planting is an order of magnitude more expensive at USD$1500-2500 per hectare^14^. Based on this, and assuming the impacts remain constant over time, we estimate the cost per additional Mg of carbon sequestered between 2025-2050 would be around USD$7 (range: USD$4 - 13) for liana cutting compared to USD$210 for enrichment planting (USD$137 - 311). The dramatic disparity between the two treatments is so large that, even if our assumptions are out by a factor of 10, the general conclusions would still hold. We note that these estimates represent only the cost of implement in the restoration treatment and exclude other management and monitoring costs.

### Liana cutting increased growth and reduced mortality

To explore the processes driving the difference between liana cutting and enrichment planting we used the repeat ALS data covering SBE. We found that 46 % of the difference was due to faster growth in the liana cutting plots, and 54 % was due to avoided tree mortality (S6). The fact that liana cutting plots experienced less disturbance than the other treatment types is clearly visible in the ALS data (less red area in Figures 2 and S3-5). To quantify this, we classified disturbance events as a decrease in canopy height of at least 5 m over a contiguous canopy area of at least 25 m^2^, which is similar to the crown area of a canopy tree in this forest^18^. Assuming each disturbance event represents the death of a single tree, then an average of 5 trees per hectare died in the liana the liana cutting plots between 2013 and 2020, compared to 10 in the enrichment planting and control plots, and 11 in the nearby primary forest at Danum Valley.

**Figure 1.**
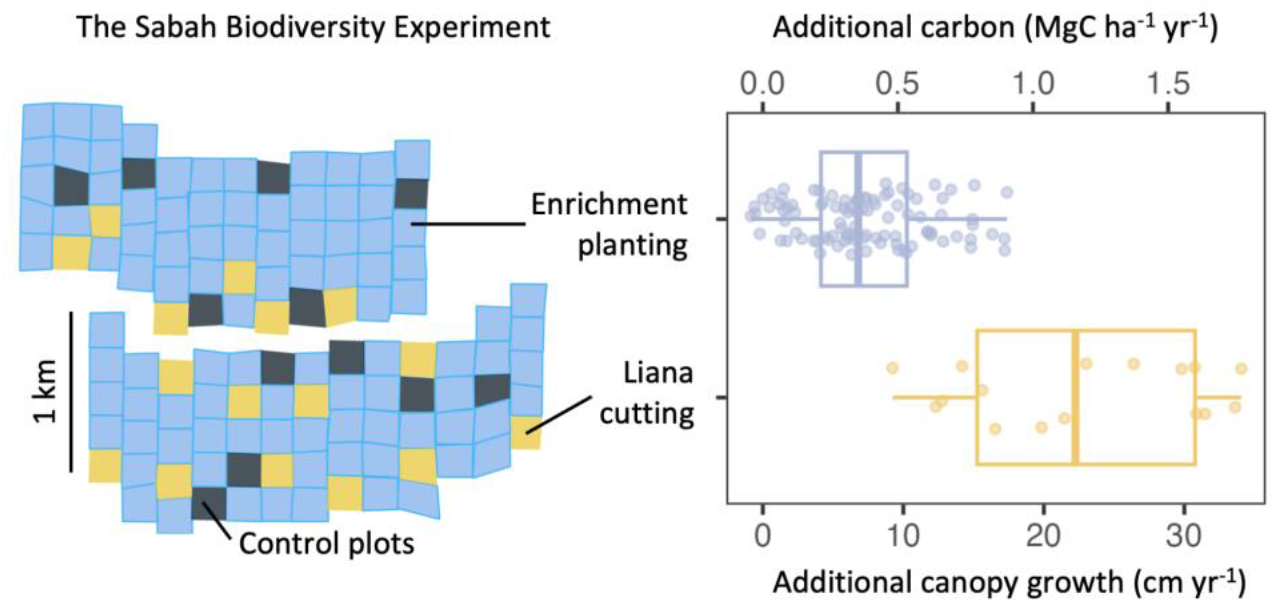
Liana cutting accelerates canopy height growth and aboveground carbon accumulation in selectively logged forests in Borneo. Left - diagram showing the layout of the Sabah Biodiversity Experiment, where each square corresponds to a 4-ha plot (n = 124). Right - Net impacts of restoration on additional canopy height growth and aboveground carbon accumulation rates. For a full comparison between experimental treatments see S1.

**Figure 2.**
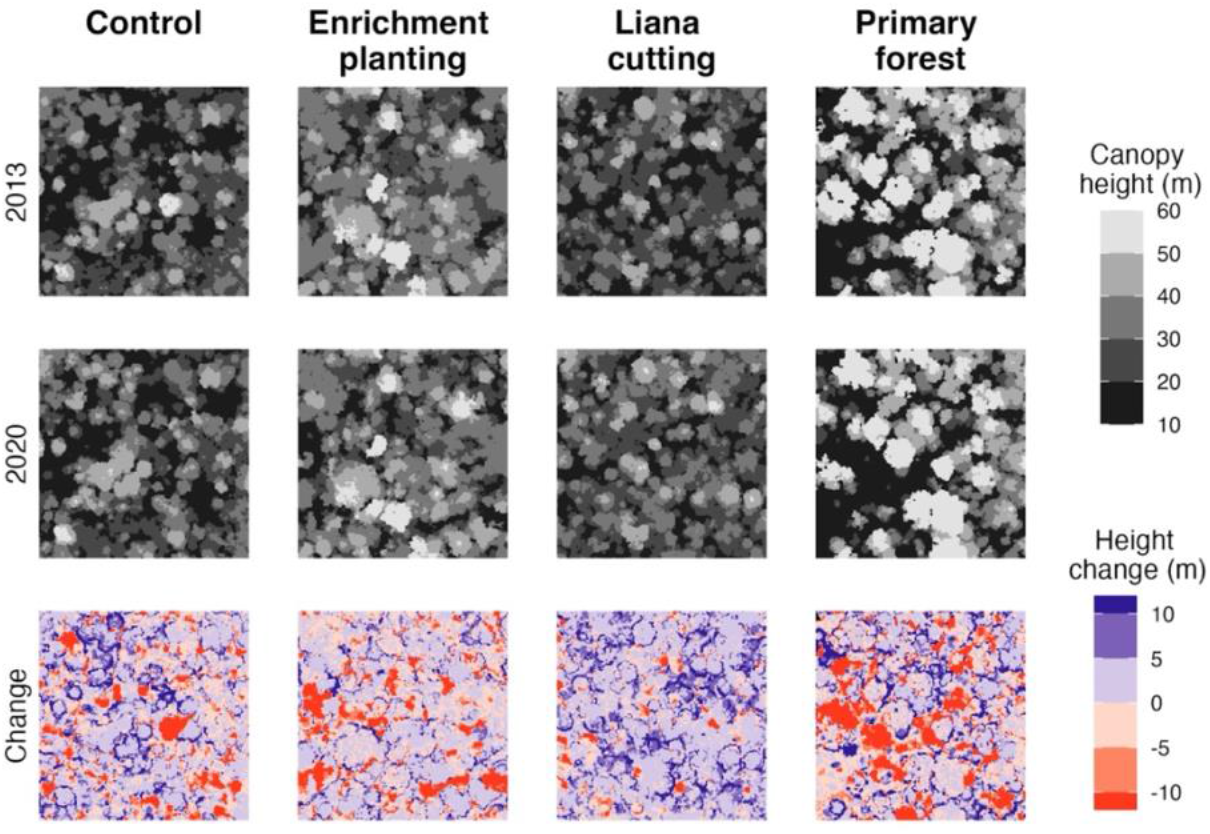
Tracking canopy dynamics using repeat airborne laser scanning. Each panel shows an example 4 ha plot. The top row shows the canopy height models (CHMs) in 2013, the second row shows 2020 and the bottom row shows the change between them. For CHMs of all plots see S3-5.

In addition, plots with liana cutting had noticeably faster height growth rates in both recovering canopy gaps and in the remaining intact canopy. The mean height growth in recovering canopy gaps was 1.1 m yr^-1^ in the liana cutting plots, compared to 0.86 m yr^-1^ in the enrichment planting plots, 0.71 m yr^-1^ in the logged controls and 0.63 m yr^-1^ in the primary forest (Figure 3b). Similarly, mean height growth rates of the intact canopy were 0.45 m yr^-1^ in the liana cutting plots, compared to 0.36-0.39 m yr^-1^ across all other treatments (Figure 3b). As the vast majority of the canopy remained intact between the two ALS surveys (90% in the liana cutting plots), these seemingly small differences in canopy height growth add up to very substantial increases in canopy volume gains across the landscape associated with liana cutting (Figure 3c).

**Figure 3.**
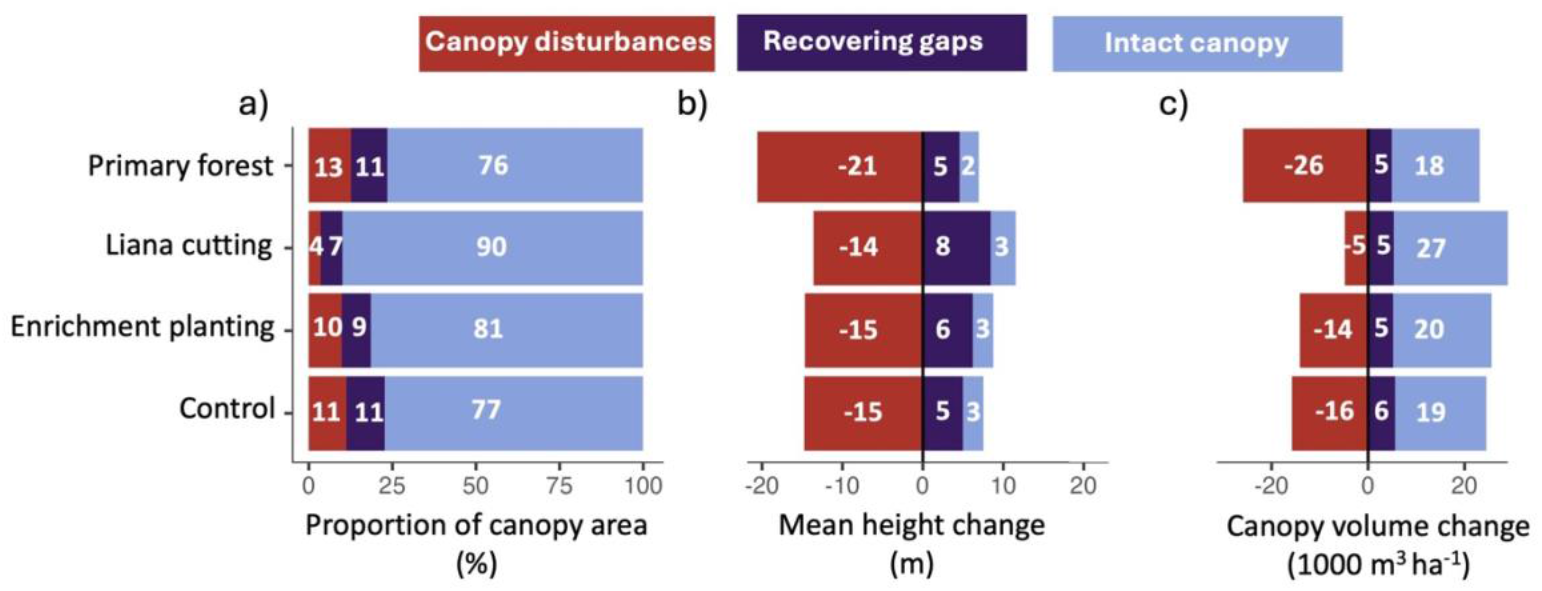
Liana cutting increased growth and reduced disturbance. Panels show (a) the area (b) the mean height change and (c) canopy volume change (area × height) in each canopy dynamics class across the experimental treatments between 2013 and 2020.

### Changing forest 3D structure after restoration

The control, enrichment planting and liana cutting plots all had a similar total leaf area index (4.13 - 4.20) and 3D distribution of leaf area in 2013 (Fig. 4a, d). As we would expect, the primary forest had a larger total leaf area index (4.40 ±0.03), concentrated at greater heights due to the presence of taller trees. Between 2013 and 2020, the leaf area of the control and enrichment plots increased in the upper-canopy (35-45 m), suggesting that the canopy trees are still growing after logging (Fig 4f). Over the same period their leaf area decreased in the mid-canopy (10-30 m), suggesting that the planted seedlings had not yet reached these heights in sufficient density to offset the effects of tree mortality we observed in Figures 2 & 3. These two effects balanced out so that the total leaf area index of the control and enrichment plots did not change substantially between 2013 and 2020. By contrast, the leaf area of the liana cutting plots increased significantly across all heights from 10-45 m, reflecting the faster growth and lower mortality described above. This resulted in an overall leaf area index of 4.53 ±0.02 in the liana cutting plots in 2020, higher even than the primary forest which decreased slightly to 4.28 ±0.03.

**Figure 4.**
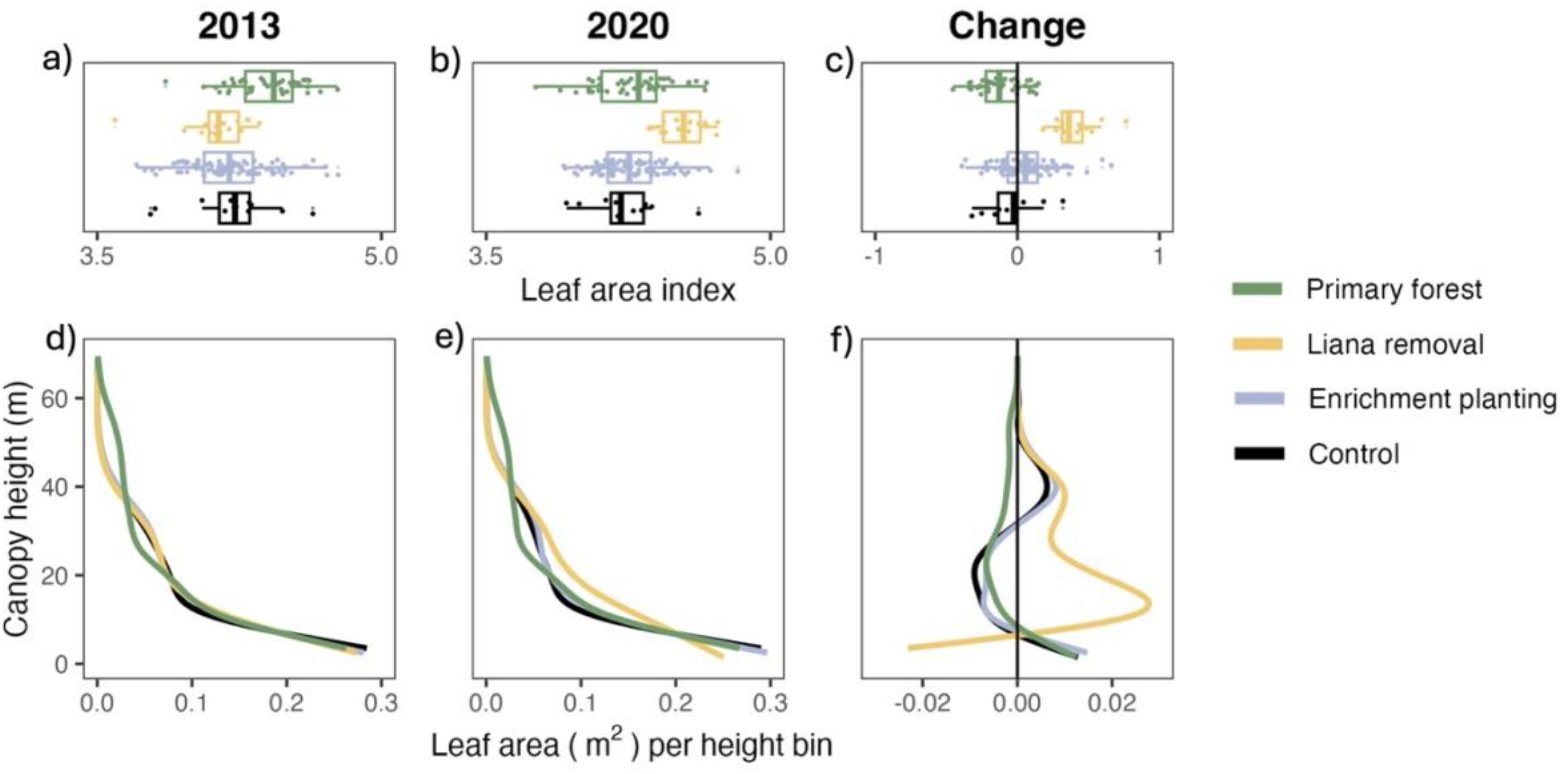
Liana cutting alters 3D forest structure. The top row (a-c) shows the variation in leaf area index for each 4-ha plot. The bottom row (d-f) shows vertical leaf area profiles of leaf area. The columns show the values in 2013 (a & d); 2020 (b & e) the difference between them (c & f).

## Discussion

Large areas of forests in Borneo were selectively logged in the 1970s and 1980s, typically at much higher intensities than those in the Amazon or Congo Basin^20^. Many of these logging concessions are no longer actively managed, which increases their risk of being cleared and replaced with oil palm^21^. Our study demonstrates that liana cutting could provide a cost-effective opportunity to restore these forests and extend management over large areas of at-risk forests that might otherwise be lost. We found that liana cutting accelerated accelerating forest growth three times faster than enrichment planting, sequestering carbon at a cost of USD$7 per ton (range: USD$4 - 13). The magnitude of this effect was unexpected given the general consensus that severe lianas infestations are much less common in the dipterocarp forests of Southeast Asia compared to other tropical regions^14–16^.Previous estimates have high uncertainty ranges and derive from biologically distinct regions^8–10^, so it is unclear whether they are transferable to the dipterocarp dominated forests of Southeast Asia. By contrast, the scale and duration of the Sabah Biodiversity Experiment means that our estimate for the cost-effectiveness of liana cutting are directly relevant to forest management in Borneo.

To understand the ecological impacts of liana cutting, we must examine the processes driving forest recovery. Multiple studies in central America have shown that liana cutting releases trees from competition, enabling them to grow faster^8,9,22^. We found a similar response in Borneo, with liana cutting leading to both greater height growth rates of mature trees that make up the intact canopy, and faster regeneration in canopy gaps compared to enrichment planting plots and logged forest controls. However, this accelerated growth only accounted for around half of the increased aboveground carbon accumulation following liana cutting. The remaining difference between restoration treatments was linked to the reduction in tree mortality we observed in the liana cutting plots. Based on the area disturbed, we estimate that liana cutting avoided the death of five trees per hectare, compared to the control plots. This reduced mortality is of a similar magnitude as the number of trees typically harvested in a round of selective logging in Borneo, which generally targets between 4-10 trees per hectare every 30-year cutting cycle^19^.

There are several mechanisms through which liana cutting could lead to reduced tree mortality^23^. Lianas can physically suppress trees and outcompete them for light^24^. Their greater hydraulic efficiency can also reduce a tree’s access to water and could exacerbate the effects of drought^25^. Our study overlapped with a severe drought in Borneo caused by the 2015-16 El Niño^26^, and it is possible that liana cutting alleviated this enough to reduce tree mortality. Beyond competition with trees, lianas can also enhance risk of tree mortality by lightning^27^, wind gusts during storms^28^ and as collateral damage during tree falls^23^. Of course, liana cutting may have simply delayed tree mortality, rather than reducing it in the long-term. Between 2013 and 2020, the canopy in the liana cutting plots at SBE became increasingly closed and dense, which may well lead to enhanced competition for light followed by self-thinning. Our results provide a snapshot of the first decade following liana cutting, but understanding the long-term implications of this restoration intervention will require continued monitoring both at SBE and across the tropics.

While it is tempting to think of lianas simply as structural parasites of trees, it is important to remember that they also play a vital ecological role for many other species – as a source of food and by facilitating movement through the canopy. For example, research in both the Neotropics and Borneo has shown that lianas generally promote greater bird abundance and diversity^34,35^. The fact that lianas adopt very different functional strategies to trees, geared towards more acquisitive leaf and root traits^25,36,37^, suggests that their removal could also have knock-on impacts on forest biogeochemical cycling^38^. While lianas typically only make up a small fraction of the aboveground biomass in a tropical forest, they contribute disproportionately to total leaf area, and therefore presumably to primary productivity^39,40^. Systematically removing lianas could therefore have unforeseen impacts on carbon and nutrient cycling in the litter and soil that have so far been almost completely unexplored. These impacts could potentially be mitigated by partial cutting of lianas (e.g., cutting 60-80% of stems as opposed to all of them), which is currently being trialed in the Ulu Segama-Malua Forest Reserve near SBE^41^. This work will hopefully shed light on how best we can achieve the structural recovery benefits of liana cutting without compromising biodiversity and ecosystem function.

## Methods

### The Sabah Biodiversity Experiment

The Sabah Biodiversity Experiment (SBE) is a forest restoration and tree biodiversity experiment in Sabah, Malaysian Borneo, that has been running since 2002^17^. The region is characterized by a tropical climate, with a mean annual temperature of around 27°C and mean annual rainfall exceeding 3000 mm. SBE covers a 500 ha area of lowland dipterocarp forest that was selectively logged in the 1980s and has since been allowed to recover (S1). The area is divided into two blocks (north and south) separated by an old logging road and is topographically heterogeneous, with elevation ranging between 150-350 m a.s.l.

The experimental design of SBE consists of 124 plots (200×200 m in size, or 4 ha), randomly allocated to different experimental treatments (64 in the southern block and 60 in the north). 12 plots were designated as controls and were allowed to recover without any post-logging intervention. All remaining plots received understory enrichment planting with varying combinations of 16 dipterocarp species. Specifically, 32 plots were planted with a single dipterocarp species, 32 plots with a mixture of four species, and a further 32 plots with all 16 species. Additionally, a further 16 plots planted with the full mixture of 16 species were also subjected to liana cutting (10 plots in the south block and 6 in the north). Due to field constraints, the 6 plots in the northern block were not randomly allocated but located close to the access road (see Fig 1) and therefore at relatively low elevation.

Seedlings were planted in parallel lines 10 m apart, with a 3-m spacing between seedlings along the lines. Initial planting occurred in 2002-2003, with a second cohort planted in 2008-2009 to replace any seedling that had died. In total, around 100,000 seedlings were planted (approximately 900 per 4-ha plot, excluding the controls). Liana cutting in the 10 plots in the southern block was undertaken in 2011 and repeated again in 2014. In the remaining 6 plots in the north block, liana cutting occurred later in 2014 and was repeated in 2017. Liana cutting targeted all climbers (including bamboos) across the entire 4-ha plot with no minimum size limit.

For the purpose of our analysis, we grouped plots into three categories: logged forest controls (12 plots, 48 ha), enrichment planting only (96 plots, 384 ha) and liana cutting with enrichment planting (16 plots, 64 ha). It is important to note that the SBE design does not fully separate the effects of liana cutting from enrichment planting, as all plots (except the controls) were planted. However, the liana cutting treatment was implemented several years after enrichment planting (2011-14), meaning that any differences between treatments should have emerged more recently. Moreover, all enrichment planting plots also received a small amount of liana cutting along the planting lines (covering approximately 10% of the plot area) prior to planting and during each subsequent census in 2011-12 and 2023-24. Consequently, we can exclude that liana cutting effects are mediated by enrichment planting (e.g., increased seedling growth and survival) as all planted seedlings were cleared of lianas.

### Airborne laser scanning acquisition and processing

Airborne laser scanning (ALS) data were collected in November 2013 and February 2020 by the survey company Ground Data Solutions using a Riegl scanner mounted on a helicopter. Both ALS surveys covered the entire SBE landscape, as well as 180 ha of primary forest at nearby Danum Valley. To enable comparisons between the two sites, we gridded the data over Danum into 4-ha plots (45 plots). Data acquisition followed similar flightlines and specifications in both surveys (S7), with the 2013 data collected from an altitude of 350 m and achieved a density of 26 pulses m^-2^, while the 2020 data were flown at 250 m and a density of 32 m^-2^. Ground points were classified using the LASground function in LAStools^42^ and the point cloud was normalized by subtracting the ground elevation from the remaining points. We then used the point cloud data to build 1-m resolution digital terrain models (DTM) and canopy height models (CHM) using a locally adaptive spikefree algorithm which is highly robust to variation in ALS pulse density in the range of our data^43^.

### Additional canopy height growth and carbon accumulation

To assess the net effect of restoration we extracted the 2020 mean canopy height from the CHM using using the *terra*^44^ package in R v4.5^45^. We then modelled canopy height as a function of elevation, treatment and experimental block using the following linear model.

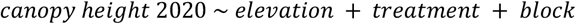

Restoration treatment was included as a factor with 3 levels (logged forest controls, enrichment planting and liana cutting) and experimental block as a factor with two levels (north and south). A table showing model results are given in S1 and post-hoc pairwise comparisons among restoration treatments were calculated using the *marginaleffects* package^46^.

To display the additional carbon accumulation due only to restoration, we corrected for the modulating effect of elevation (S1) by predicting the 2020 canopy height of each plot as if it was located at the site mean elevation (217 m). The additional canopy height growth due to enrichment planting was then calculated as the difference between enrichment planting and control plots (identical to the coefficient of the enrichment planting term in the model output). The additional canopy height growth due to liana cutting was calculated as the difference between the mean canopy height of liana cutting and enrichment planting plots. This is because liana cutting happened in addition to enrichment planting. Finally, because these different treatments started at different times we divided the additional canopy height in 2020 by the period of the treatment (18 years for enrichment planting, 9 years for liana cutting in the southern block and 6 years for liana cutting in the northern block). This gave the additional canopy height growth shown in Fig 1.

To convert the additional canopy height growth into additional carbon accumulation we used a simple allometric equation developed specifically for Sabah^12^. We found that both linear and power law models gave very similar carbon estimates (S2) so chose to use the linear model for simplicity. This linear model simply predicts that each additional metre of growth in mean canopy height equates to 5.2 Mg Carbon per hectare (see S2).

To compare the cost effectiveness of liana cutting and enrichment planting we wanted to estimate the cost of sequestering a ton of carbon by 2050, assuming the intervention takes place now (2025). To make this calculation, we first assumed that the treatment effect similar over 25 years. This seems justified for the enrichment planting since we already have 18 years of data, but is a larger extrapolation from the 9-years of liana cutting. We then combined the 25 year carbon accumulation estimates with the cost of the restoration treatment in SBE. These costs were derived from the contracts used to implement the restoration and ranged from USD$140-330 per hectare for the liana cutting and USD$1500-2500 per hectare for the enrichment planting. We took the central value from these ranges and also calculated the best and worst case scenarios by combining the lowest cost with the highest effectiveness and vice-versa.

### Disturbance and recovery dynamics

To disentangle the processes driving canopy dynamics, we used a recently developed framework^47^ that classifies repeat ALS data into three classes of canopy dynamics: new disturbances, recovering canopy gaps, and intact canopy. Disturbances were defined as contiguous areas larger than 25 m^2^ which decreased in height by more than 5 m between scans. Recovering gaps were defined as contiguous areas larger than 25 m^2^ which had a canopy height lower than 10 m in the initial 2013 scan. Intact canopy is the remaining area of the CHM that was not classified as a recovering gap in 2013 and was not subjected to a disturbance between the two scans. This classification makes full use of the repeat ALS data and helps tease apart the processes driving forest recovery, such as tree mortality, understory growth in gaps, and mature tree growth. For each 4-ha plot at SBE and Danum, we calculated the proportion of the canopy classified as either disturbed, recovering gap or intact, and then quantified the mean canopy height change in each of these three classes. We then multiplied these together to get the change in canopy volume over time attributed to new disturbances, recovering gaps and intact canopy growth.

The differences between liana cutting and enrichment planting were due to a combination of faster growth in the intact canopy and gaps, as well as a lower area of canopy disturbed (avoided mortality). To estimate the relative contributions of these two factors to the overall difference between the treatments, we built a counterfactual treatment which had the same area of intact canopy, gaps and disturbance as the enrichment planting. We then multiplied these areas by the height change rates of the liana cutting plots, giving a counterfactual volume change due to increased growth only (S6). We then converted this volume change into a change in mean canopy height by dividing by the plot area, and compared mean canopy height increases between the enrichment planting, liana cutting and counterfactual treatments.

### Vertical leaf area index profiles

To explore the impact of restoration on different canopy layers, we used the *leafR* package^48^ to generate vertical LAI canopy profiles of each SBE plot in 1-m height bins. Leaf area profiles were generated from the normalized point clouds by estimating the transmittance of laser pulses through the canopy. Since ALS samples the canopy from above, the lower levels of the canopy are likely to be occluded. The *leafR* package therefore applies a Beer-Lambert transformation to the transmittance profiles to estimate leaf area in each 1-m height bin. This transformation assumes homogeneous volume filling through the canopy, which is rarely the case in a forest. However, as we were not interested in the absolute values of LAI, but rather the difference between treatments, this approach allowed us to compare the vertical canopy profiles of enrichment planted and liana cutting plots and pinpoint where they differ in 3D space.

Specifically, for each restoration treatment we calculated the mean vertical LAI profile across all plots. We repeated this calculation for both the 2013 ALS scan, which occurred at around the same time as the liana cutting and therefore likely too early to detect any differences between treatments, and the 2020 scan which took place 6-9 years after liana cutting. Additionally, we also calculated the total LAI of each plot in 2013 and 2020 by summing the leaf area across all height layers.

## Data and code availability

The airborne laser scanning data collected for this study is available on Zenodo: https://zenodo.org/records/14917551

The scripts to reproduce the analysis are available on Github: https://github.com/TobyDJackson/Repeat_LiDAR_liana_cutting

## Acknowledgments

We thank the Danum Valley Management Committee and Sabah Biodiversity Council for access and assistance with the airborne laser scanning surveys in Malaysia (permit number JKM/MBS.1000-2/2 JLD.9 122). We also thank Ground Data Solutions, who collected all the airborne laser scanning data used in this study. TJa, AH, EJ, DAC, DFRPB and TJu were supported by a NERC Standard Grant (NE/X000281/1). TJu was additionally funded by a NERC Independent Research Fellowship (NE/S01537X/1) and a Research Project Grant from the Leverhulme Trust that also supported FJF (RPG-2020-341). The 2013 airborne laser scanning data collection was funded by ETH Zurich. The 2020 airborne laser scanning data collection was funded through a NERC Standard Grant awarded to DAC (NE/S010750/1) and a NERC Independent Research Fellowship awarded to TJu (NE/S01537X/1). For the purpose of open access, the author has applied a Creative Commons Attribution (CC BY) licence to any Author Accepted Manuscript version arising from this submission.

