## Supplementary materials for "Liana cutting accelerates tropical forest recovery at a fraction of the cost of tree planting"

#### S1. Mean canopy height growth by treatment and block

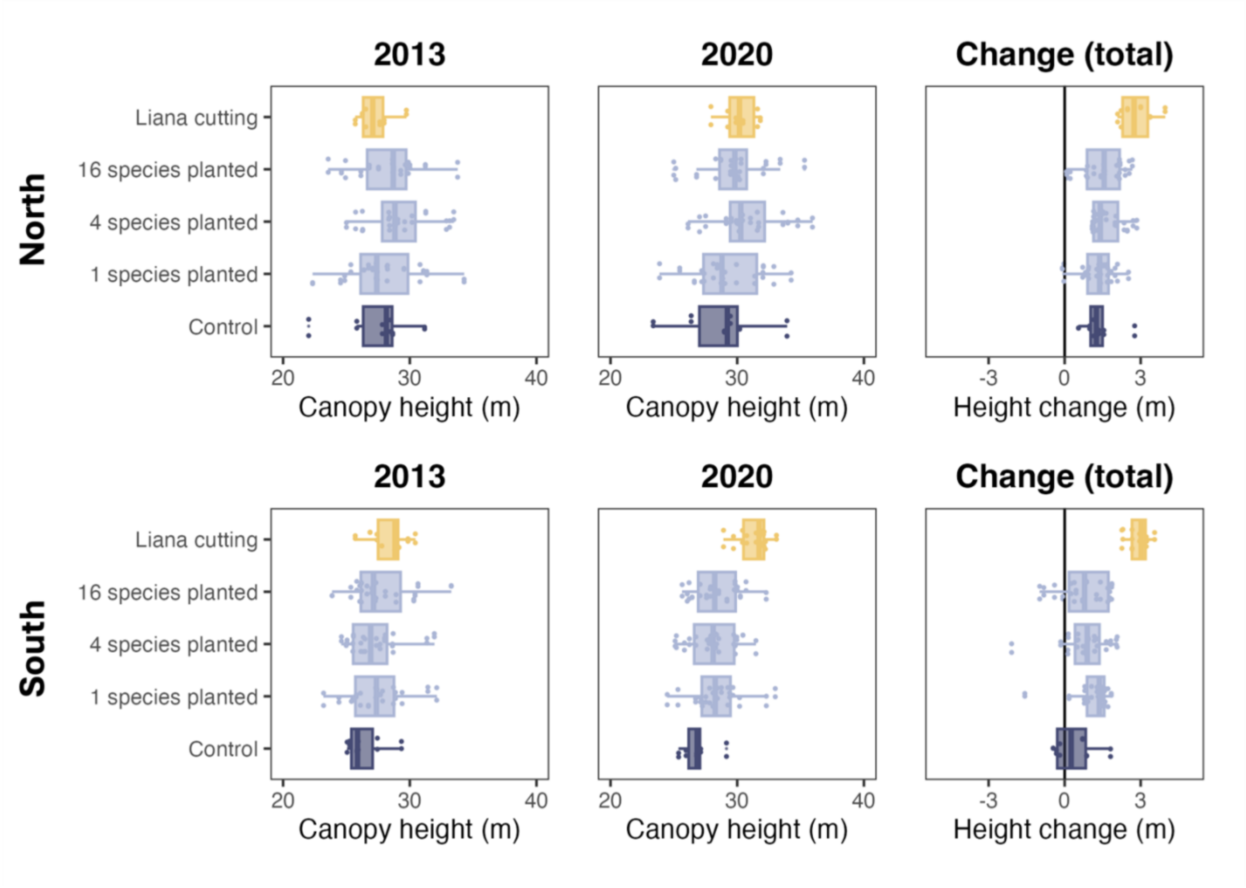


***Fig. S1***: *Variation in mean canopy height mapped from ALS data in 2013 (left), 2020 (centre) and the change between the two periods (right) across all experimental treatments. The top row shows the plots in the northern block, while the bottom row shows the plots in the southern block. Results shown in the central panels are analogous to those presented in Fig. 1 of the main text, where the 1, 4 and 16-species mixture treatments (light blue boxplots) were pooled.*

### S2. Topographic impact on canopy height and recovery

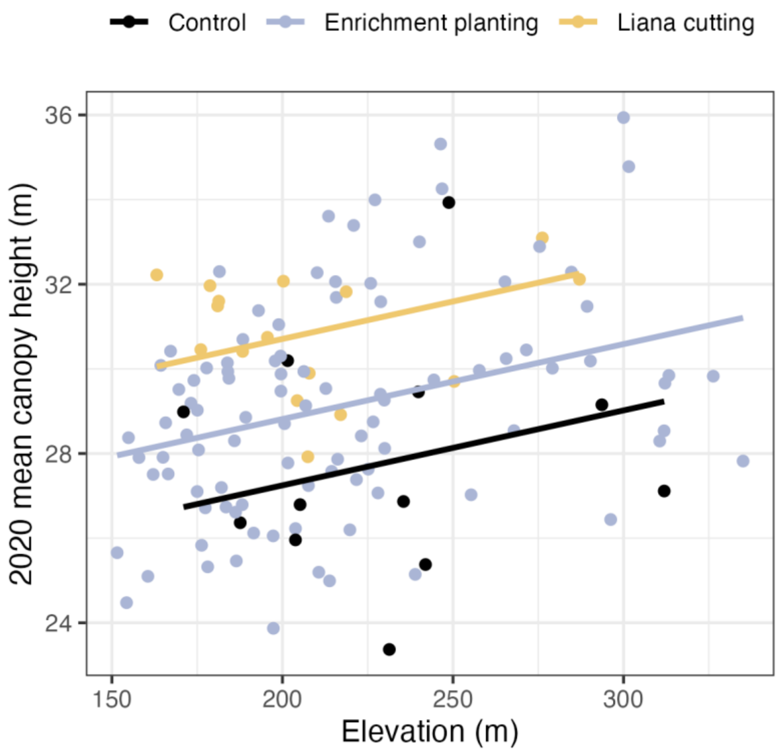


*Figure S2 - 2020 mean canopy height against elevation (m.a.s.l) for each 4 ha plot in the Sabah Biodiversity Experiment. The lines show the linear model with treatment modelled as a factor with three levels.*

| **term** | **estimate** | **std.error** | **p.value** |
|---|---|---|---|
| Intercept | 25.4 | 1.0 | 0.00 |
| Elevation | 0.0 | 0.0 | 0.00 |
| Control | -1.6 | 0.7 | 0.02 |
| Liana cutting | 2.1 | 0.6 | 0.00 |
| Southern block | -1.6 | 0.4 | 0.00 |

*Table S2 - Output of the linear model for 2020 mean canopy height.*

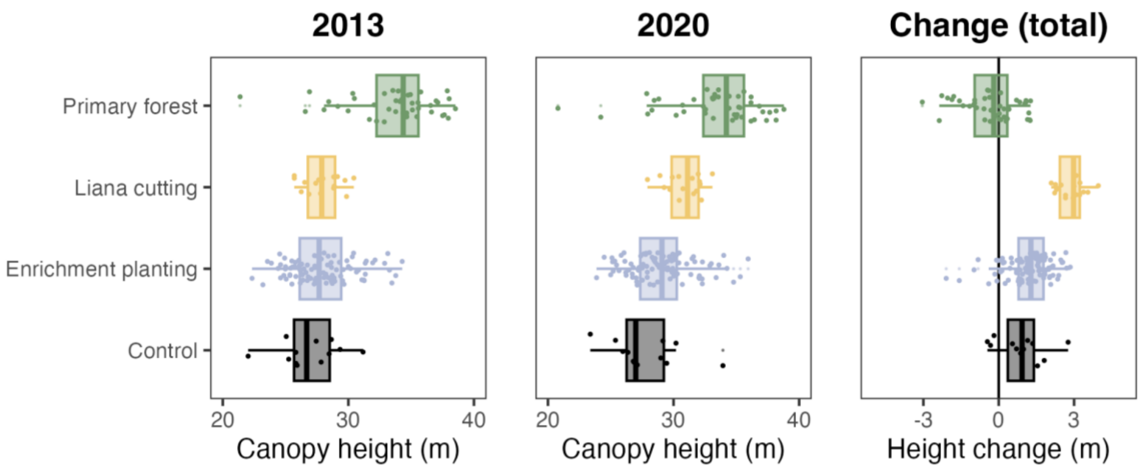


*Figure S1.2 - Boxplots showing mean canopy height in 2013, 2020 and the change between them for each treatment in the Sabah Biodiversity Experiment as well as the nearby primary forest at Danum.*

### S3. Carbon accumulation due to treatments

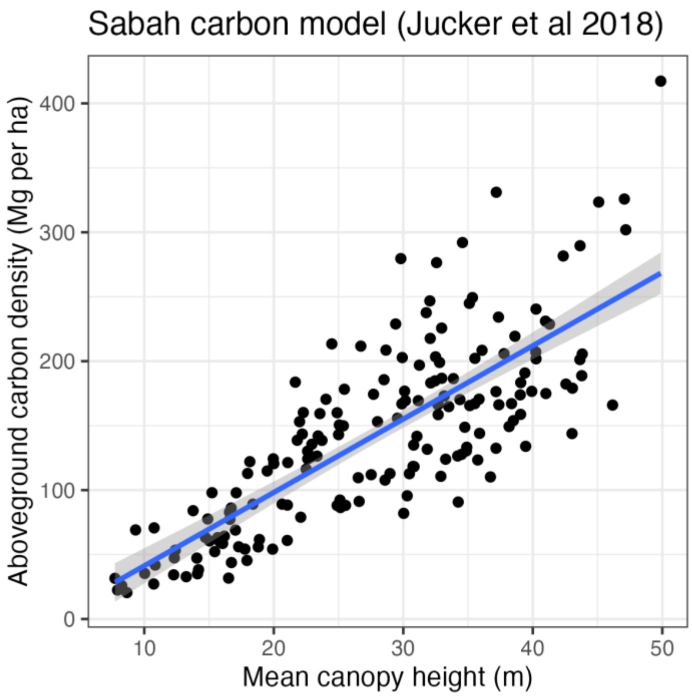


*Figure S2 - Data from Jucker et al (2018) showing above ground carbon density against mean canopy height for Sabah forests. This data was used to develop the allometric equations to predict carbon storage for the plots in the Sabah Biodiversity Experiment.*

In the main text we used a linear model to estimate carbon storage from mean canopy height. Using this approach we estimated that liana cutting accumulated +1.31 $\pm$ 0.13 Mg C $ha^{-1}$ $yr^{-1}$, three times greater than that of enrichment planting (+0.38 $\pm$ 0.03 Mg C $ha^{-1}$ $yr^{-1}$). We also tested using a power law allometric model, which estimated that liana cutting accumulated +1.53 $\pm$ 0.15 Mg C $ha^{-1}$ $yr^{-1}$ and enrichment planting accumulated +0.44 $\pm$ 0.03 Mg C $ha^{-1}$ $yr^{-1}$.

| | Enrichment planting (2002 - 2020) | Liana cutting (2013 - 2020) |
|---|---|---|
| (A) Additional canopy height growth (m) | 1.6 0.7 | 1.9 0.3 |
| (B) Additional carbon sequestration (Mg $ha^{-1}$) | 9.2 3.9 | 10.7 1.7 |
| (C) Experiment period (years) | 18 | 6.25 |
| (D) Carbon accumulation (Mg $ha^{-1}$ $yr^{-1}$) | 0.5 0.2 | 1.7 0.3 |
| (E) Restoration cost (USD$ $ha^{-1}$) | 1500 - 2500 | 140 - 330 |
| (F) Cost of carbon to 2050 (USD$ $Mg^{-1}$) | 157 (64 - 1269) | 5 (2.5 - 11.2) |

*Table 1: Cost-effectiveness of liana cutting and enrichment planting in terms of medium-term carbon sequestration (present day to 2050). The cost of carbon (expressed as Mg, where 1 Mg = 1 tonne or 1000 kg) to 2050 is calculated as F = E/(25×D). The range in A, B and D are standard errors. The range of costs in E are from the contracts used in Sabah and from the literature. The range in F represents the variation from a situation where the lowest cost (E) is combined with the highest effectiveness (D) of each treatment, to one where the highest cost occurs with the lowest effectiveness. We note that these costs represent implementation costs only and do not include the cost of verification or other management processes.*

#### S4. Height change rasters - liana cutting

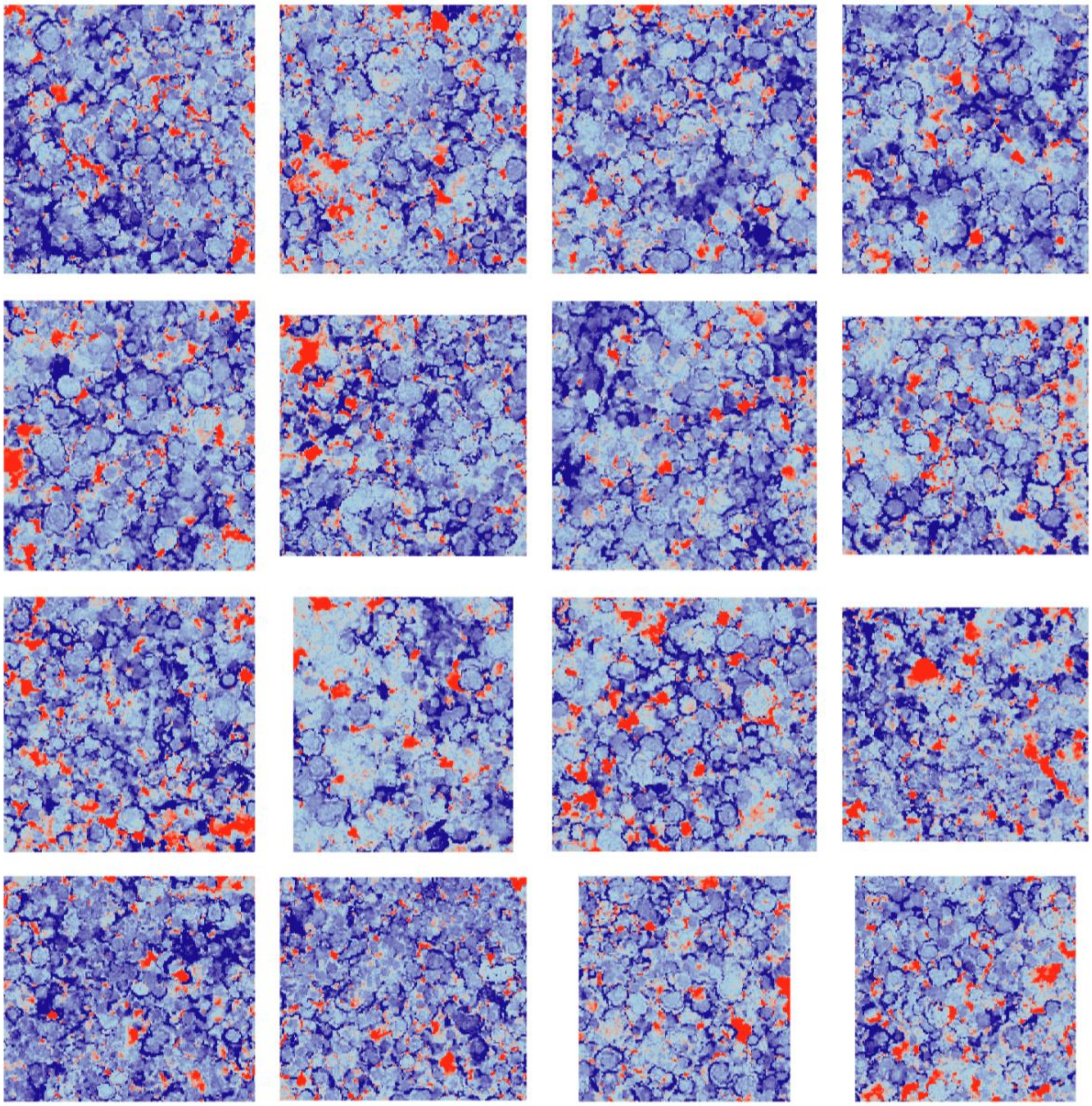

*Figure S3 - Height change rasters for all liana cutting plots.*

#### S5. Height change rasters - enrichment planting (16 species)

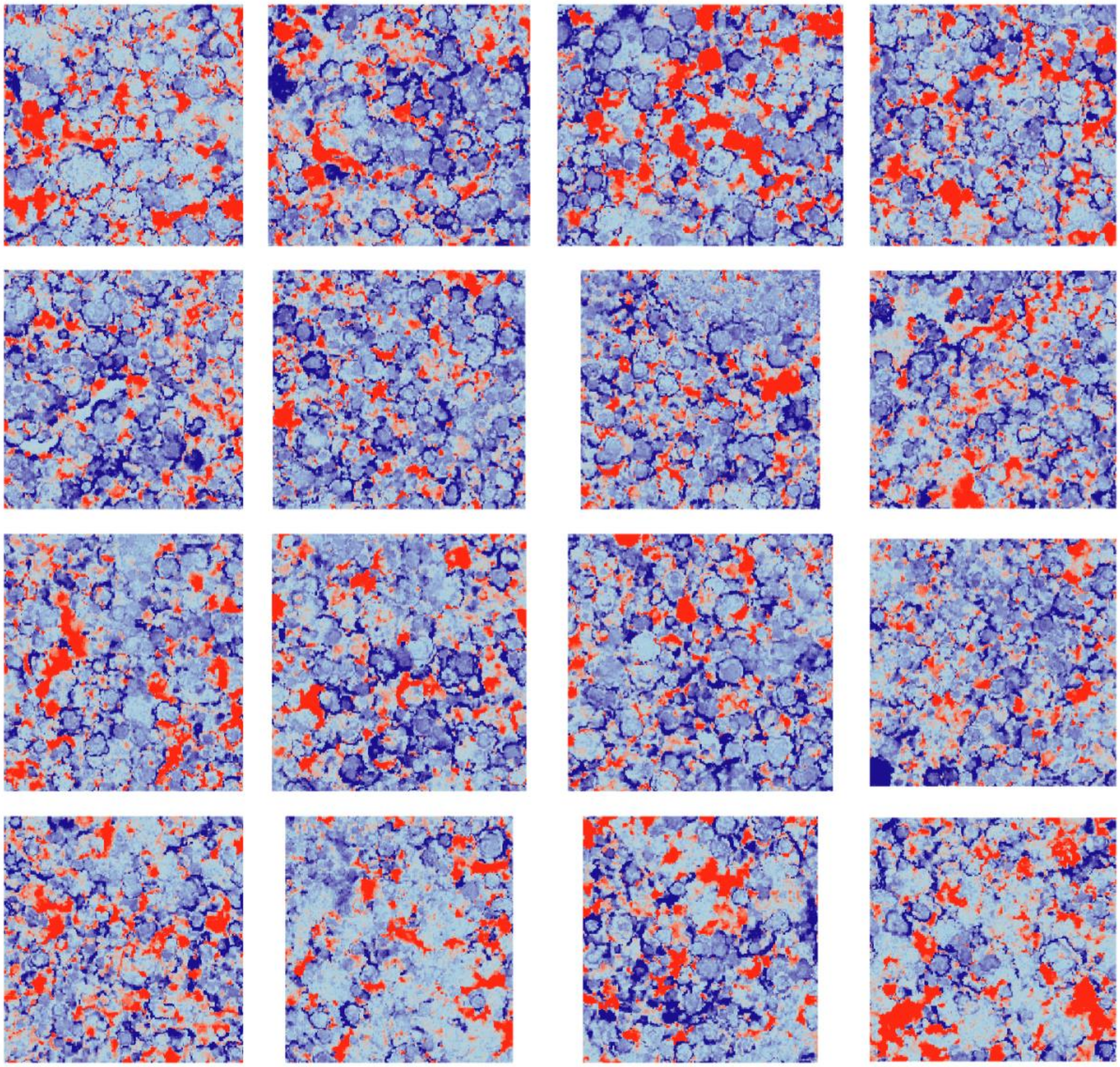

*Figure S4 - Height change rasters for all 16-species enrichment planting plots which did not undergo liana cutting. These are the most directly comparable to the liana cutting plots.*

#### S6. Height change rasters - control plots

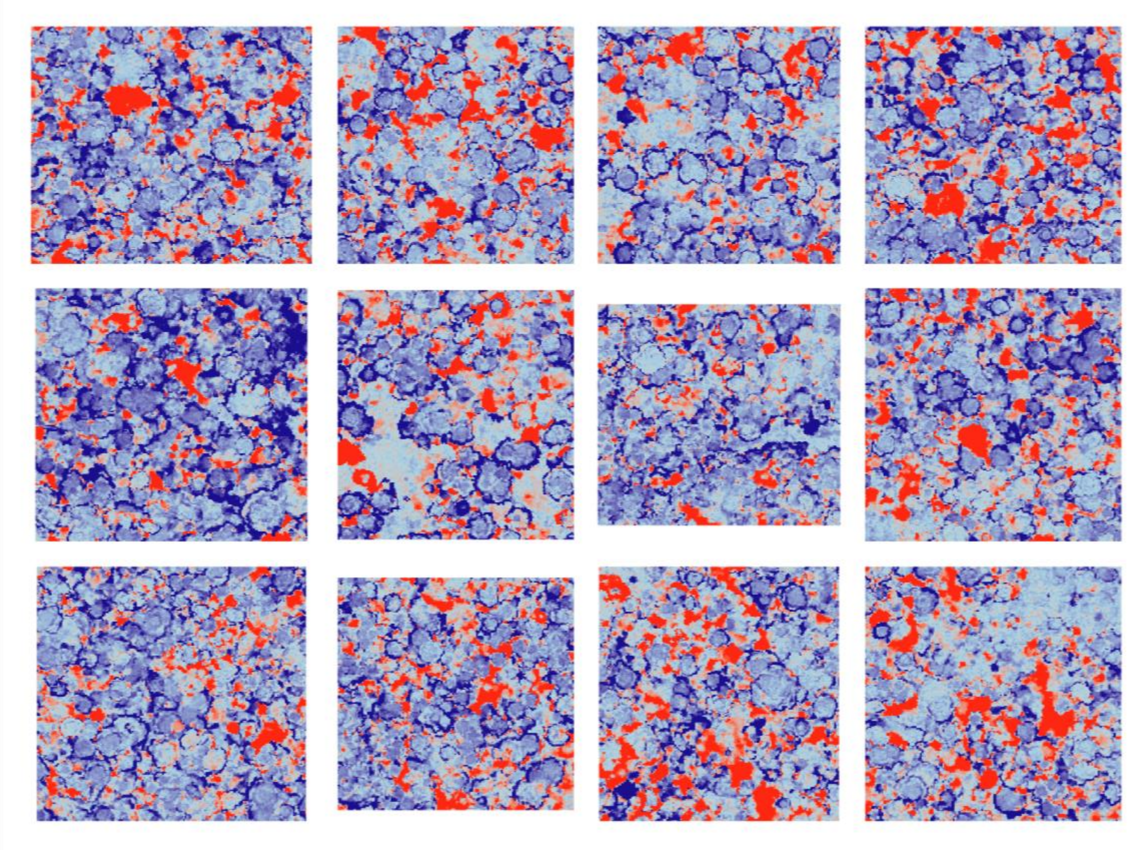

*Figure S5 - Height change rasters for all control plots.*

#### S7. Counterfactual approach to estimating relative importance of faster growth and avoided mortality after liana cutting

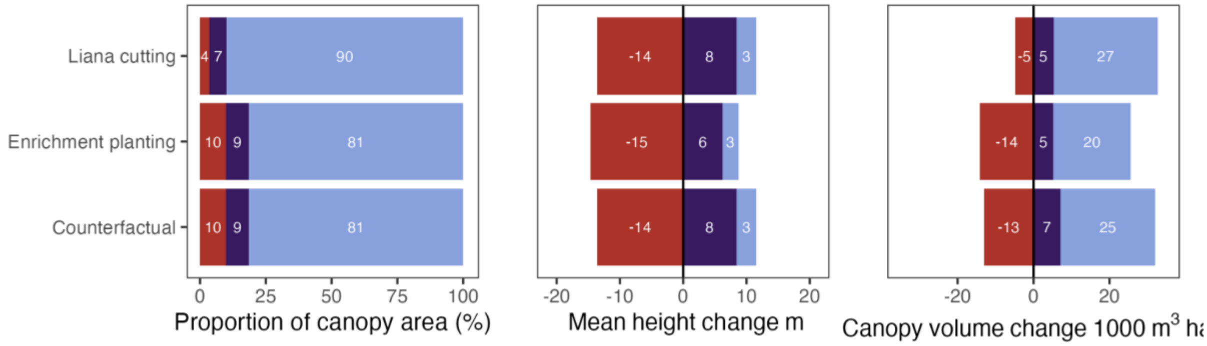


*Figure S6 - Similar to Fig 3 shows the area, height change and canopy volume change across the three canopy dynamics classes. This figure compares liana cutting, enrichment planting and a counterfactual which has the same proportion of canopy area in each canopy dynamics classes*

*as the enrichment planting, but the same mean height change in each class as the liana cutting. This counterfactual enables us to calculate*

| | Mean height increase (m) |
|---|---|
| (A) Enrichment planting | 1.15 |
| (B) Liana cutting | 2.78 |
| (C) Counterfactual | 1.90 |
| (D) Liana cutting - enrichment planting (B-A)<br>(this is due to both faster growth and lower disturbance) | 1.63 |
| (E) Counterfactual - enrichment planting (C-A)<br>(this is only due to faster growth) | 0.75 |

*Table S6. Calculation of the relative contribution of avoided disturbance and accelerated growth to the change in mean canopy height in the liana cutting plots. Note that the height increases reported here were calculated without correcting for elevation and so are slightly different from those reported in the main text.*

The proportion of growth due only to faster growth can therefore be calculated as the ratio of E to D, which is 46%.

#### S8. Airborne laser scanning details

| | 2013 survey | 2020 survey |
|---|---|---|
| Date | 17/11/2013 | 19/02/2020 |
| Day of year | 321 | 50 |
| Scanner type | Riegl | Riegl |
| Altitude | 200 | 250 |

| | | |
|---|---|---|
| Beam divergence (mrad) | 0.5 | 0.5 |
| Pulse density | 26 | 32 |
| Max scan angle (°) | 10 | 10 |

*Table S7. Details of airborne laser scanning data used in this study.*

| Scenario | N trees per hectare | | Basal area per hectare | |
|---|---|---|---|---|
| | Dipterocarp | Other | Dipterocarp | Other |
| Unlogged (Danum 50 ha data) | 196 | 4939 | 11.5 | 18.3 |
| Logged (remove 500 biggest dipts) | 186 | 4939 | 3.0 | 18.3 |
| Liana cutting (all trees grow by 10%) | 186 | 4939 | 3.6 | 22.1 |

*Table S7. Details of airborne laser scanning data used in this study.*
By comparison, the latest seedling census gives a total basal area of 0.1 m^2 per hectare.
